# Golgi localized β1-adrenergic receptors stimulate Golgi PI4P hydrolysis by PLCε to regulate cardiac hypertrophy

**DOI:** 10.1101/640789

**Authors:** Craig A. Nash, Wenhui Wei, Roshanak Irannejad, Alan V. Smrcka

## Abstract

Increased adrenergic tone resulting from cardiovascular stress leads to development of heart failure, in part, through chronic stimulation of β1 adrenergic receptors (βARs) on cardiac myocytes. Blocking these receptors is part of the basis for β-blocker therapy for heart failure. Recent data demonstrate that G protein-coupled receptors (GPCRs), including βARs, are activated intracellularly, although the biological significance is unclear. Here we investigated the functional role of Golgi βARs in cardiac myocytes and found they activate Golgi localized, prohypertrophic, phosphoinositide hydrolysis, that is not accessed by cell surface βAR stimulation. This pathway is accessed by the physiological neurotransmitter norepinephrine (NE) via an Oct3 organic cation transporter. Blockade of Oct3 or specific blockade of Golgi resident β1ARs prevents NE dependent cardiac myocyte hypertrophy. This clearly defines a physiological pathway activated by internal GPCRs in a biologically relevant cell type and has implications for development of more efficacious β-blocker therapies.

## Introduction

Cardiovascular disease, particularly heart failure, is the leading cause of disease and death in the developed world. In response to long term pathological stress, such as hypertension or myocardial infarction, the levels of neurohumoral factors such as epinephrine, endothelin and angiotensin II increase. These hormones directly stimulate G protein-coupled receptors (GPCRs), including β-adrenergic receptors in cardiac myocytes (1–3). Chronic stimulation of these receptors, including stimulation of β-adrenergic receptors (βARs) by catecholamines, drives cardiac hypertrophy and ventricular remodeling, ultimately leading to heart failure. The efficacy of β-blocker therapy for treatment of heart failure results, at least in part, by ameliorating chronic βAR stimulation in the heart (4, 5).

Previous studies by our group, and others, have implicated specific phospholipase C (PLC) enzymes downstream of Gq and Gs-coupled GPCRs in the regulation of cardiac hypertrophy. PLCβ isoforms stimulated by Gq have been implicated in cardiac hypertrophy driven by cell surface a-adrenergic receptors (aAR) (6, 7). Our laboratory identified a critical role of PLCε in cardiac hypertrophy driven downstream of the endothelin (ET-1A) receptor and cAMP signaling by βARs (8, 9). PLCε is poised as a nexus for multiple receptor signaling systems due its diversity of upstream regulators including heterotrimeric G protein βγ subunits and small GTPases, including Rap, Rho and Ras, and cAMP via the Rap guanine nucleotide exchange factor (GEF), Epac (10–14).

PLCε and Epac are scaffolded together at the nuclear envelope in cardiac myocytes by the hypertrophic organizer, muscle-specific A kinase anchoring protein (mAKAPβ), in close proximity to the Golgi apparatus (15). Activation of PLCε in cardiac myocytes induces hydrolysis of phosphatidylinositol-4-phosphate (PI4P) at the Golgi apparatus leading to release of diacylglycerol (DAG) and inactive IP_2_ in the vicinity of the nucleus to facilitate nuclear protein kinase D (PKD) and subsequent downstream hypertrophic pathways (9, 16). In previous work we found that either the Epac-selective cAMP analog, cpTOME, or forskolin stimulation of adenylate cyclase, activate PLCε-dependent Golgi PI4P hydrolysis in cardiac myocytes. Surprisingly, the βAR agonist isoproterenol (Iso), does not stimulate Golgi PI4P hydrolysis despite strongly stimulating cAMP production (8, 9). As an explanation for this apparent paradox we demonstrated PLCε-dependent PI4P hydrolysis can be controlled by two distinct pools of cAMP delimited by distinct PDE isoforms. One pool, limited by PDE3 acts through Epac and Rap1 to activate PLCε, while a second pool controlled by PDE2 and/or PDE9A inhibits PLCε activity through activation of PKA (17). cAMP generated downstream of Iso cannot access the Epac/mAKAPβ/PLCε complex unless PDE3 is specifically inhibited partially explaining the lack of activation of Golgi PI4P hydrolysis by Iso.

A new paradigm for generating localized cAMP signals in cells has been established by a number of recent studies indicating that GPCRs in different intracellular compartments have discrete signaling outputs (18–22). An emergent idea is that βARs internalized into endosomes continue to stimulate cAMP accumulation resulting in cellular outcomes distinct from those generated by βARs at the cell surface. Irannejad *et al* established that β1ARs expressed at the Golgi apparatus in HeLa cells can induce G protein activation in the Golgi when stimulated by either a cell permeant agonist, dobutamine, through passive diffusion, or the physiological hormone epinephrine, when transported across the cell membrane by the organic cation transporter, Oct3 (19). These ideas led us to hypothesize that in cardiac myocytes, βARs resident at the Golgi apparatus could generate cAMP that has privileged access to the Epac/mAKAPβ/PLCε scaffold at the nuclear envelope/Golgi interface, thereby yielding a set of signals divergent from those generated by cell surface βARs. We demonstrate that endogenous β1ARs at the Golgi apparatus in cardiac myocytes are required to stimulate hypertrophic Epac/PLCε-dependent PI4P hydrolysis at the Golgi, and that these intracellular βARs can be accessed by physiological neurotransmitters, and synthetic β-blockers and agonists. These data present a potential new paradigm for drug development for treating heart failure through deliberate targeting of internal βARs in cardiac myocytes.

## Results

### Dobutamine induces β1-AR activation and PI4P hydrolysis at the Golgi

In previous studies we showed that cAMP produced upon stimulation with the βAR agonist, Iso, does not induce PI4P hydrolysis at the Golgi in NRVMs without addition of a PDE3 inhibitor (17). We hypothesized that intracellular βARs may be required to generate a specific pool of cAMP with privileged access to the Epac/mAKAPβ/PLCε complex, and that the cell impermeant agonist, Iso, is unable to access these internal receptors. To test this hypothesis, we utilized the membrane permeant β1AR-agonist, dobutamine, either alone, or in combination with the βAR antagonists, metoprolol and sotalol, which are membrane-permeant and -impermeant, respectively (19). NRVMs were transduced with an adenovirus expressing the PI4P biosensor, FAPP-PH-GFP and stimulated with dobutamine (100 nM) or PBS. In contrast to Iso, dobutamine stimulated rapid and sustained PI4P hydrolysis (Fig. 1A, left). Addition of metoprolol, but not sotalol, blocked dobutamine-stimulated PI4P hydrolysis (Fig. 1A middle and right panels). This indicates that stimulation of an intracellular population βARs is required for dobutamine-mediated PI4P hydrolysis. To visualize βAR activation at the Golgi apparatus, NRVMs transfected with translocatable Venus-based sensor of Gs coupled receptor activation, NES-Venus-mini-Gs (23). When a Gs coupled receptor is activated, NES-Venus-mini-Gs binds to the activated receptor, which is then visualized as translocation from the cytoplasm to membranes where the activated receptor is located providing information on receptor activation in subcellular compartments. In unstimulated cells NES-Venus-mini-Gs is distributed throughout the cytoplasm. We did not detect translocation of fluorescence to endogenously expressed receptors (data not shown), so NRVMs were cotransfected with the β1AR. Addition of dobutamine to cells co-expressing NES-Venus-mini-Gs and β1AR, caused a rapid clearance of cytoplasmic fluorescence and accumulated fluorescence at the PM and punctate structures likely corresponding to nascent t-tubules. This was followed by slower translocation of Venus associated fluorescence to the perinuclear region of the cell colocalizing with the CFP-giantin, a marker for the Golgi apparatus (Fig. 1B, D and E and Movie S1). Iso also caused rapid translocation to the PM and t-tubules but, in contrast to dobutamine, no accumulation at the perinuclear region was observed (Fig. 1C, D and E and Movie S2). To further confirm association of NES-Venus-mini-Gs with the Golgi, cells were treated with dobutamine for 8 min to cause NES-Venus-mini-Gs translocation to the perinuclear region, followed by treatment with Brefeldin A. Brefeldin A treatment significantly reversed NES-Venus-mini-Gs association with the perinuclear region, confirming Golgi localization of NES-Venus-mini-Gs (Fig. S1 and Movie S3). Taken together, these data support the idea that a population of βARs present at the Golgi can be activated by exogenous agonists and stimulate Gs, ultimately resulting in PLC activation and PI4P hydrolysis.

**Figure 1.**
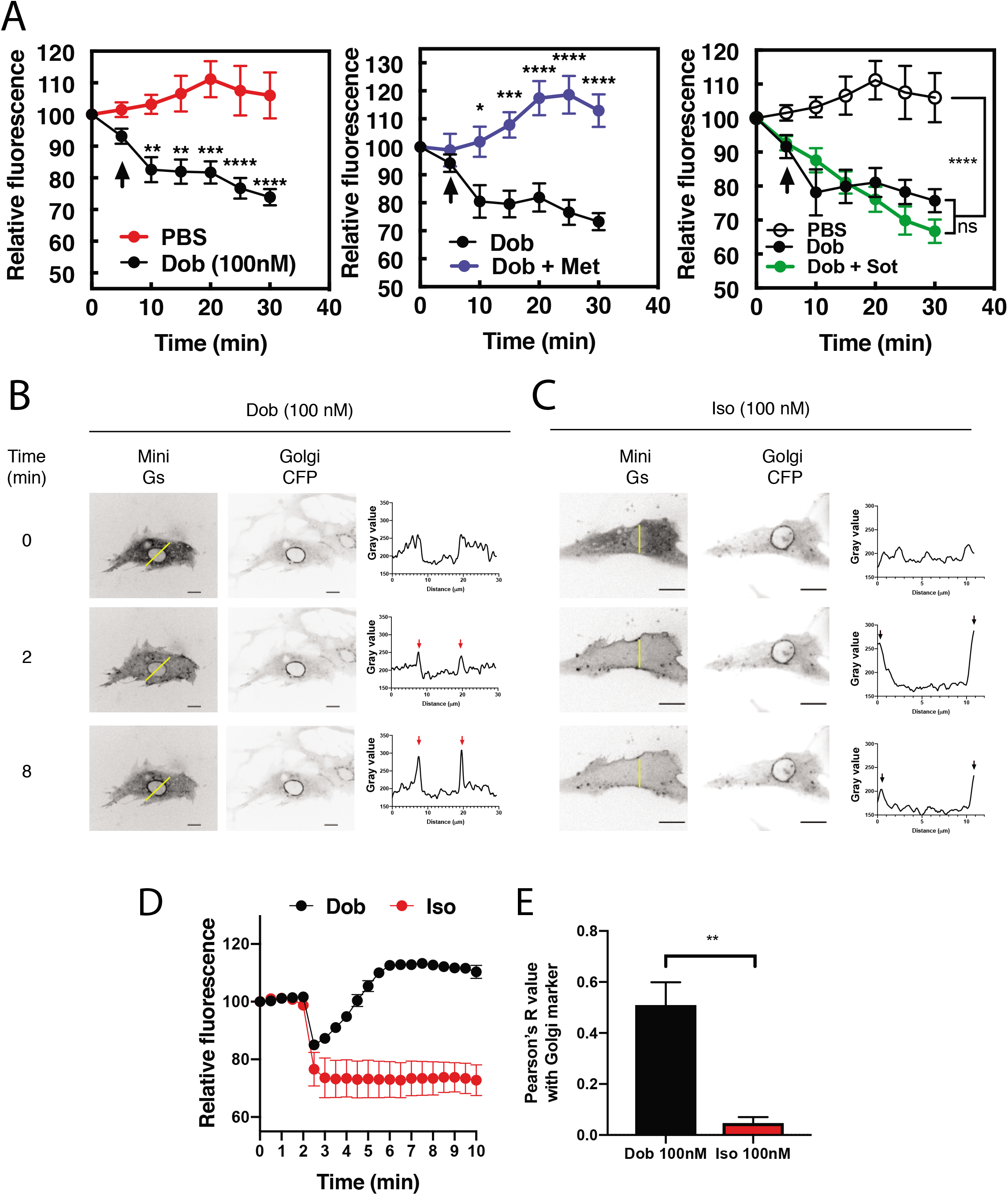
Dobutamine induces PI4P hydrolysis through the activation of internal βARs. A) NRVMs were transduced with FAPP-PH-GFP and stimulated as indicated. Time lapse live cell microscopy was used to quantitate Golgi associated FAPP-PH-GFP fluorescence was quantitated as previously described (9, 11). NRVMs were stimulated with dobutamine alone (100 nM, left), dobutamine in the presence or absence of metoprolol (100 μM, center), or dobutamine in the presence or absence of sotalol (5 mM, right). Data are not significantly different between dobutamine and dobutamine + sotalol. Data are from at least n=12 cells each from 3 separate preparations of NRVMs. B) NRVMs were transfected with β1-ARs and NES-Venus-mini-Gs, followed by viral transduction with CFP-Giantin. Representative images of dobutamine-mediated NES-Venus-mini-Gs recruitment (100 nM, B left), CFP-Giantin Golgi marker (B, center) and histogram of representative NES-Venus-mini-Gs recruitment (B, right). Red arrow = Perinuclear region, Black arrow = sarcolemma. The yellow line indicates where histogram data was captured. Scale bars are 10 μm. C) Representative images of Iso-mediated NES-Venus-mini-Gs recruitment (100 nM, left), CFP-Giantin Golgi marker (B, center) and histogram of representative NES-Venus-mini-Gs recruitment. Yellow line indicates where histogram data was captured. Scale bars=10 μm D) Mean data of fluorescence intensity of NES-Venus-mini-Gs at the perinuclear region corresponding to the Golgi +/− SEM from at least 5 cells. E) Pearson’s correlation coefficient for overlap of YFP-Mini-Gs and CFP-Giantin images. All time course graphs are presented as mean ± standard error. Agonists were added where as indicated by the arrow.

### PI4P hydrolysis by dobutamine requires Golgi-localized PLCε and Epac

Previous data from our laboratory demonstrated that the Epac/mAKAPβ/PLCε complex is responsible for cAMP-mediated PI4P hydrolysis at the Golgi in NRVMs (9, 17). To determine if this complex is responsible for dobutamine stimulated PI4P hydrolysis we utilized adenoviral shRNA to deplete PLCε, or adenoviral transduction of the RA1 domain of PLCε, which competes for the interaction between PLCε and mAKAPβ, disrupting the mAKAPβ-PLCε complex (8). PLCε shRNA depletion of PLCε completely inhibited PI4P hydrolysis stimulated by dobutamine, while control scrambled shRNA had no effect (Fig. 2A). Viral expression of the PLCε RA1 domain also inhibited PI4P hydrolysis stimulated by dobutamine (Fig. 2B). This demonstrates that mAKAPβ-scaffolded PLCε is required for dobutamine-mediated PI4P hydrolysis. The Epac-selective inhibitor HJC0726 inhibited PI4P hydrolysis stimulated by dobutamine (Fig. 2C), however, the βγ inhibitor Gallein had no effect (Fig. 2D). Taken together, these data indicate that Golgi PI4P hydrolysis stimulated by intracellular GPCRs requires the Epac/mAKAPβ/PLCε complex.

**Figure 2.**
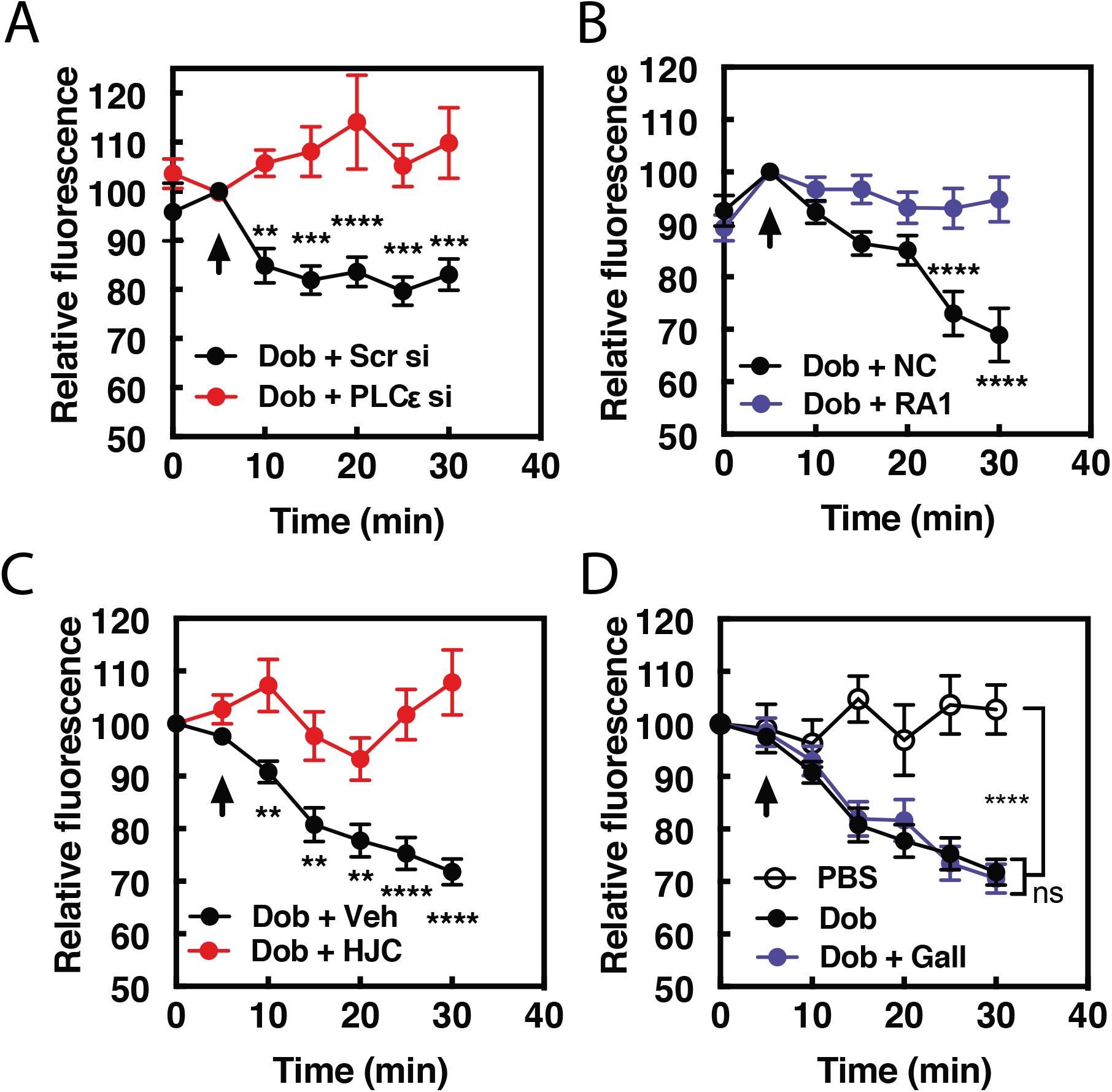
Dobutamine-mediated PI4P hydrolysis requires Epac and mAKAPβ bound PLCε. NRVMs were transduced with FAPP-PH-GFP, stimulated with treatments as indicated and Golgi associate fluorescence was monitored with time. A) PLCε knockdown prevents dobutamine-mediated PI4P hydrolysis. NRVMs were transduced for 48 hours with adenovirus expressing shRNA for PLCε or scrambled control shRNA before stimulation with 100 nM dobutamine (at arrow). B) Disruption of PLCε-mAKAPβ interaction prevents PI4P hydrolysis stimulated by dobutamine. NRVMs were transduced with RA1 domain expressing adenovirus or control virus as previously described 24 hours before experimentation. Cells were stimulated with 100 nM dobutamine (at arrow). C) Epac is required for dob-mediated PI4P hydrolysis. The Epac inhibitor HJC0726 (1 μM) was added to NRVMs 15 minutes before imaging and dobutamine (100 nM) was added at the arrow. D) Gβγ is not required for dobutamine-mediated PI4P hydrolysis. The Gβγ inhibitor, Gallein (10 μM) was added 15 minutes prior to imaging and dobutamine added as indicated by the arrow. Data are not significant between Dob and Dob + Gallein. Images for PI4P hydrolysis were taken from at least n=13 cells each from at least 3 separate preparations of NRVMs.

### Inhibition of Golgi βARs prevents PI4P hydrolysis stimulated by dobutamine

The data presented so far suggest that dobutamine acts through βARs at the Golgi apparatus to stimulate Golgi PI4P hydrolysis. To more directly demonstrate a requirement for endogenous βAR activation in the Golgi we targeted the βAR selective nanobody, Nb80, to the Golgi apparatus using the rapamycin inducible FRB-FKBP system. This approach has been previously used to inhibit βARs at the Golgi in Hela cells (19). NRVMs were transduced with virus containing CFP-Nb80-FRB and FKBP-mApple-GalT protein for Golgi targeting along with a virus expressing GFP-FAPP-PH. Cells were selected that were positive for mApple and CFP (Fig. 3A), and GFP fluorescence. In cells pretreated with vehicle control, dobutamine stimulated PI4P hydrolysis. However, following addition of 1μM Rapamycin, dobutamine-induced PI4P hydrolysis was significantly inhibited (Fig. 3B). These data confirm that βARs expressed directly on the Golgi are required for dobutamine-mediated PI4P hydrolysis at the Golgi.

**Figure 3.**
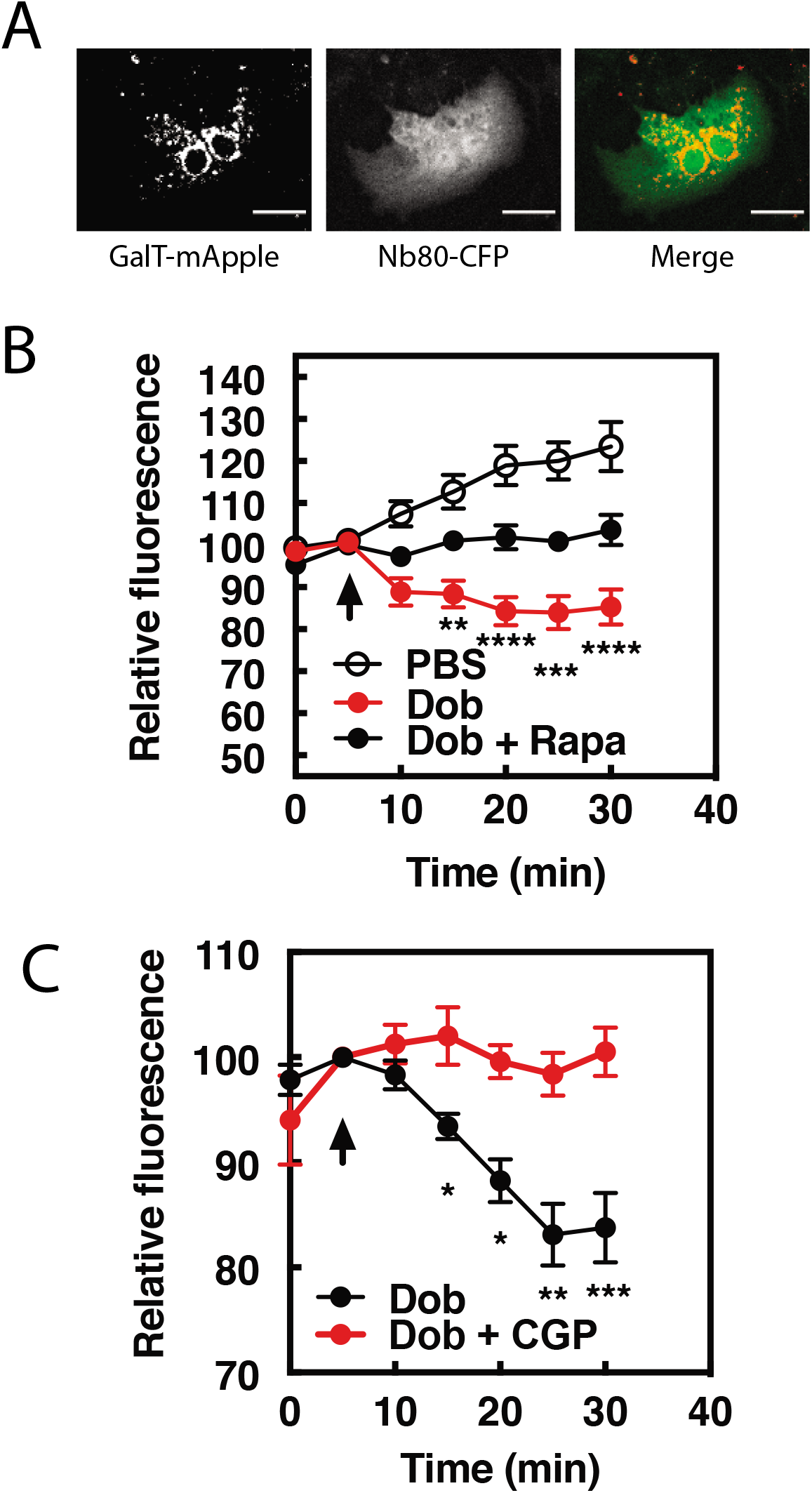
β1ARs in the Golgi are required for PI4P hydrolysis to dobutamine. NRVMs were transduced with FRB-CFP-Nb80 and FKBP-GalT-mApple containing adenovirus, along with adenovirus containing FAPP-PH-GFP for 24 hours. A) Confocal fluorescence images of NRVMs expressing FRB-CFP-Nb80 in the CFP channel and FKBP-GalT-mApple in the red channel (see methods for details). B) NRVMs were incubated with either rapamycin (1 μM) or DMSO control for 15 minutes prior to addition of dobutamine (100 nM) added at the arrow. Golgi associated FAPP-PH-GFP fluorescence was monitored by time lapse fluorescence video microscopy as in Fig. 1 A. C) Cells were pretreated treated with the cell permeable β1AR selective antagonist, CGP-20712 (100nM), or vehicle, followed by dobutamine addition and assessed as in B. Images were taken from at least n=9 cells each from at least 4 separate preparations of NRVMs. Agonists were added as indicated by the arrow.

Nb80 can bind to and block both β1 and β2ARs and while dobutamine is relatively selective for β1ARs over β2ARs, it can still potentially stimulate β2ARs. To confirm that dobutamine stimulated PI4P hydrolysis is through β1ARs, cells were treated with the cell permeant highly selective β1AR antagonist CGP-20172 (100nM) followed by stimulation with dobutamine. CGP-20172 completely blocked dobutamine-stimulated PI4P hydrolysis (Fig. 3C), indicating that the dobutamine stimulation of Golgi localized PI4P hydrolysis is through Golgi localized β1ARs.

### Norepinephrine (NE) induces PI4P hydrolysis and βAR activation at internal membranes in cardiac myocytes

Thus far we have shown that the synthetic β1AR selective agonist, dobutamine can enter cells, activate intracellular βARs, and stimulate intracellular PLC activity which has interesting pharmacological implications but may not be relevant to physiological cardiac regulation. For this reason, we determined if the sympathetic neurotransmitter norepinephrine (NE) could stimulate PI4P hydrolysis, and βAR activation at the Golgi in NRVMs. Cells transduced with adenovirus containing FAPP-PH-GFP were stimulated with NE (10 μM) at 37°C. NE caused delayed, but robust and sustained PI4P hydrolysis (Fig. 4A, left). PI4P hydrolysis initiated by NE was blocked by metoprolol (100 μM, Fig. 4A, center) but not sotalol (5 mM, Fig. 4A, right) indicating that NE can stimulate internal βARs. To further determine if NE could enter cells and activate βARs at the Golgi, we monitored NES-Venus-mini-Gs recruitment to the Golgi in response to NE. NE initially stimulated NES-Venus-mini-Gs translocation to the plasma membrane as was observed for dobutamine, followed by slower, NES-Venus-mini-Gs translocation to the Golgi (Fig. 4B and Movie S4). Thus, NE enters cardiac myocytes, accesses internal βARs, and stimulates PI4P hydrolysis.

**Figure 4.**
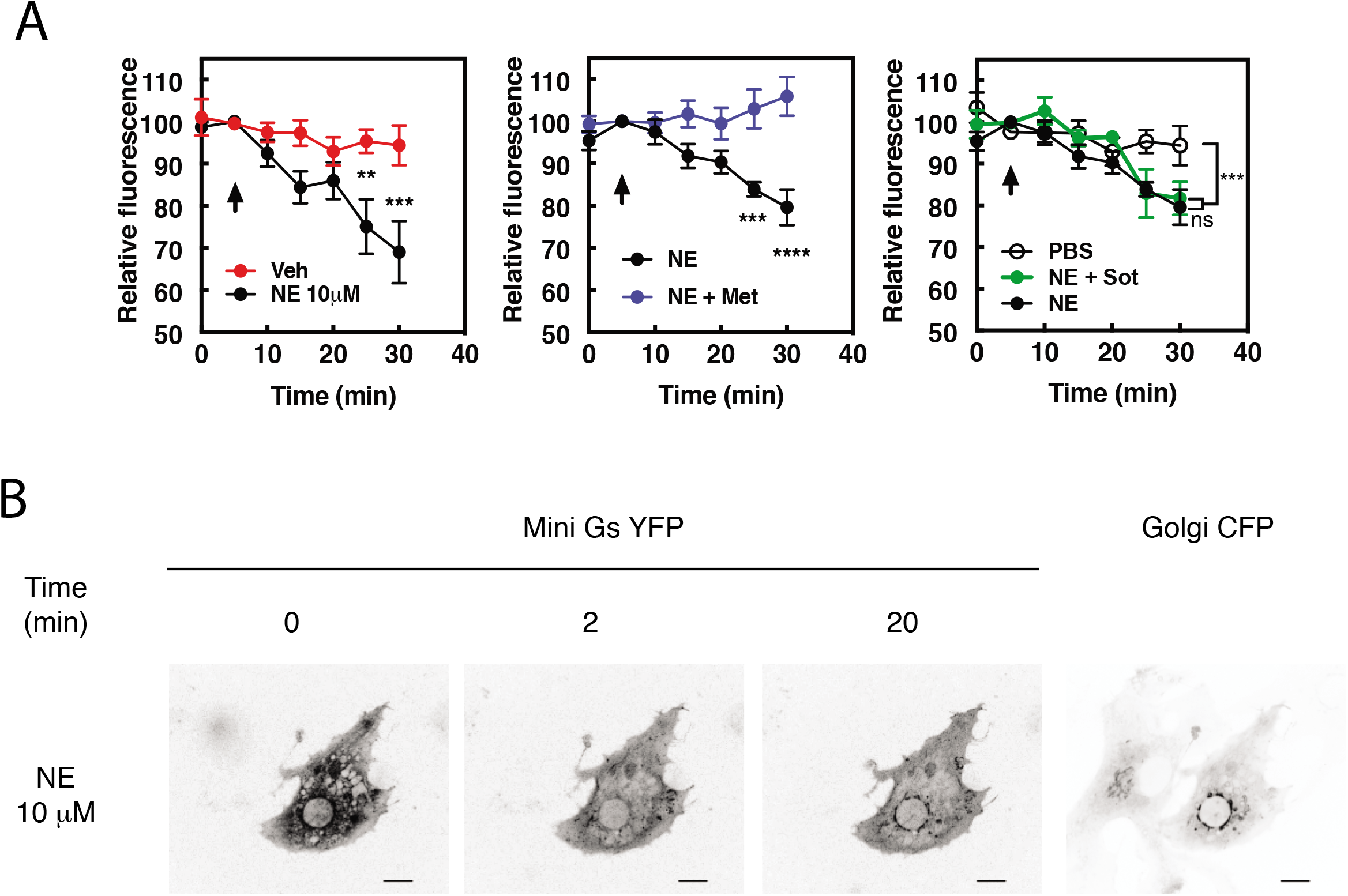
The physiological neurotransmitter, norepinephrine, can induce PI4P hydrolysis through internal receptors and can activate βARs at the Golgi. A) NRVMs were transduced with FAPP-PH-GFP and stimulated with norepinephrine (10 μM, left) in the presence of metoprolol (100 μM, center) or sotalol (5 mM, right) and analyzed as in Fig. 1A. Data are not significant between norepinephrine and norepinephrine + sotalol. B) NRVMs were transfected with β1AR and NES-Venus-mini-Gs, followed by viral transduction with CFP-Giantin. Representative images of norepinephrine-mediated NES-Venus-mini-Gs recruitment (10μM, left columns), CFP-Giantin Golgi marker (B, right). All experiments were performed in humidified environmental chamber at 37°C. Images for PI4P hydrolysis collected as in Fig. 1A, were from at least n=7 cells each from 3 separate preparations of NRVMs. Agonists were added where indicated by the arrow.

### Inhibition of the membrane cation transporter, OCT3, prevents norepinephrine induced PI4P hydrolysis

Data from other laboratories has demonstrated that transport of norepinephrine or epinephrine into cells by organic cation transporter family of proteins (OCT proteins) is required for activation of internal receptors in adult cardiac myocytes (24) and HELA cells (19). The non-selective cation transporter, OCT3 has been shown to be responsible for this transport, therefore we utilized three structurally distinct inhibitors of OCT3, corticosterone (100 μM, Fig. 5A), abacavir (10 μM, Fig. 5B) and lamotrigine (10 μM, Fig. 5C) (25) to determine if OCT3 is required for NE-stimulated intracellular PI4P hydrolysis. Preincubation of NRVMs with any of these inhibitors prevented NE from inducing PI4P hydrolysis, suggesting that OCT3-mediated transport is required for PI4P hydrolysis by this catecholamine. In addition, we sought to determine if receptor internalization is required for NE-mediated PI4P hydrolysis and βAR activation at the Golgi. An inhibitor of dynamin-mediated receptor internalization, Dyngo (40 μM), had no effect on either PI4P hydrolysis (Fig. 5D) or recruitment of NES-Venus-mini-Gs to the Golgi membrane (Fig. 5E and Movie S5) by norepinephrine. Corticosterone (100μM) had no effect on dobutamine-mediated PI4P hydrolysis (data not shown). These data indicate that agonist transport by OCT transporters, not receptor internalization, is required for NE-mediated Golgi βAR activation and PI4P hydrolysis.

**Figure 5.**
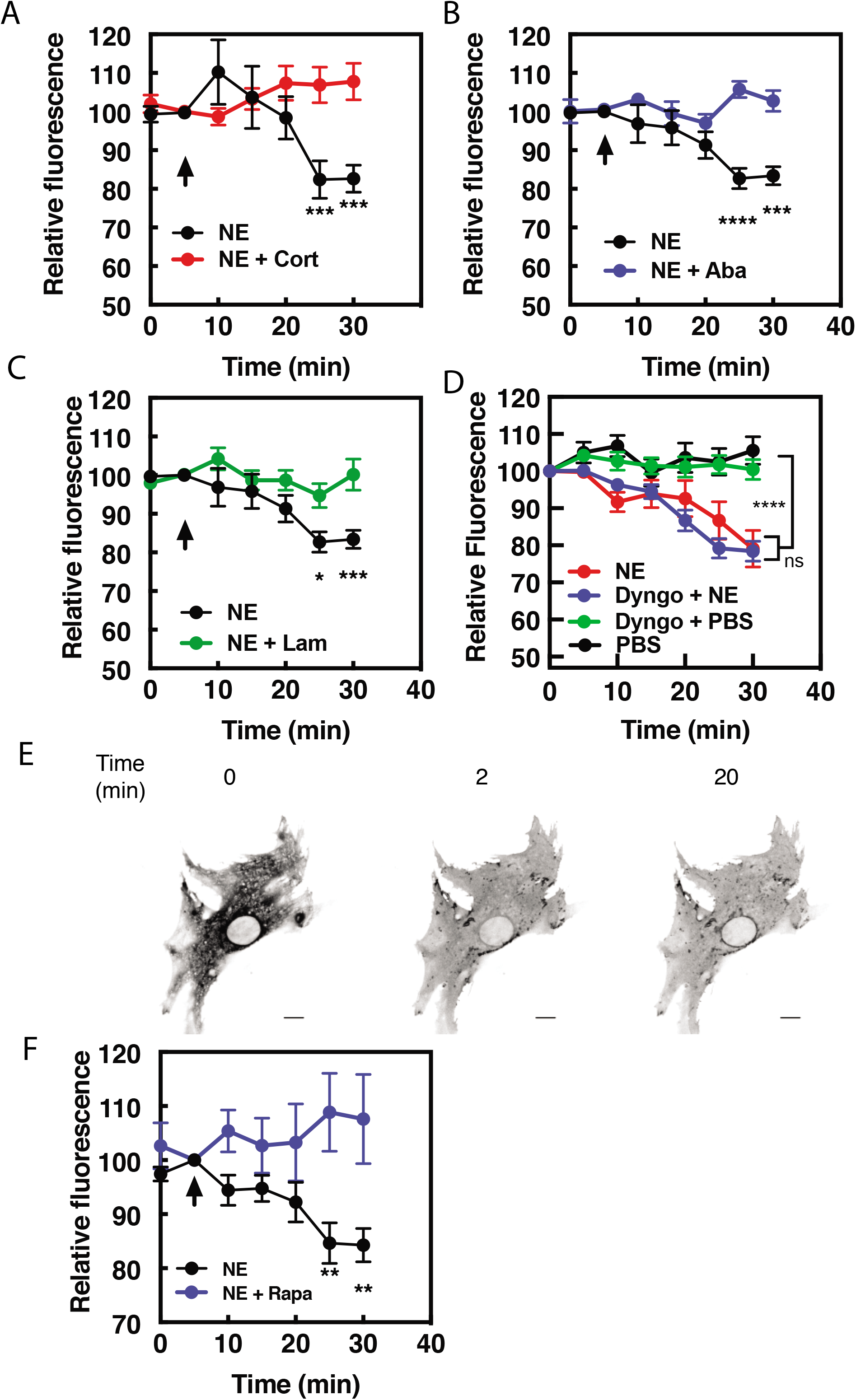
Norepinephrine requires OCT transporters but not receptor internalization to stimulated PI4P hydrolysis. NRVMs were transduced with FAPP-PH-GFP and stimulated with norepinephrine (10 μM) in the presence of corticosterone (100 μM, A), abacavir (10 μM, B), lamotrigine (10 μM, C) or Dyngo (40 μM, D). Data are not significant between norepinephrine and norepinephrine + Dyngo. Images were collected as in Fig. 1A for PI4P hydrolysis and were from at least n=10 cells each from 3 separate preparations of NRVMs. Agonists were added where indicated by the arrow. E) Dyngo has no effect on NES-Venus-mini-Gs Golgi recruitment by norepinephrine. Representative image of norepinephrine-mediated NES-Venus-mini-Gs recruitment in the presence of Dyngo (40 μM). F) NRVMs were transduced with adenovirus containing FRB-CFP-Nb80 and FKBP-GalT-mApple along with adenovirus containing FAPP-PH-GFP for 24 hours prior to experimentation. NRVMs were incubated with either rapamycin (1 μM) or DMSO control for 15 minutes prior to addition of NE (10 μM) added at the arrow. Images were collected as in Fig. 1A for PI4P hydrolysis and were from at least n=10 cells each from 3 separate preparations of NRVMs.

Additionally, NRVMs were transduced with viruses containing FRB-CFP-Nb80 and FKBP-mApple-GalT, along with FAPP-PH-GFP to determine if Golgi β1AR were required for PI4P hydrolysis to NE. In cells pretreated with vehicle, norepinephrine induces PI4P hydrolysis. However, following addition of 1μM Rapamycin, norepinephrine-induced PI4P hydrolysis was significantly inhibited (Fig. 5F).

To confirm that dobutamine and NE stimulated Golgi PI4P hydrolysis is not unique to neonatal myocytes we tested the ability of dobutamine and NE to stimulate PI4P hydrolysis in acutely isolated murine adult ventricular myocytes. Infection of AVMs for 24h with FAPP-PH-GFP labels the Golgi surrounding the nucleus as well as other structures within the myocyte as we have previously reported (Fig. 6A) (11). Stimulation of AVMs with dobutamine (Fig. 6B) or NE (Fig. 6C) stimulated PI4P depletion in the Golgi. NE stimulated PI4P hydrolysis was blocked by preincubation with the Oct3 inhibitor abacavir (Fig. 6C) indicating that NE transport into AVMs is required for stimulation of PI4P hydrolysis, as was seen in NRVMs. These data indicate that internal receptors are required for stimulation of PI4P hydrolysis in AVMs as well as NRVMs.

**Figure 6.**
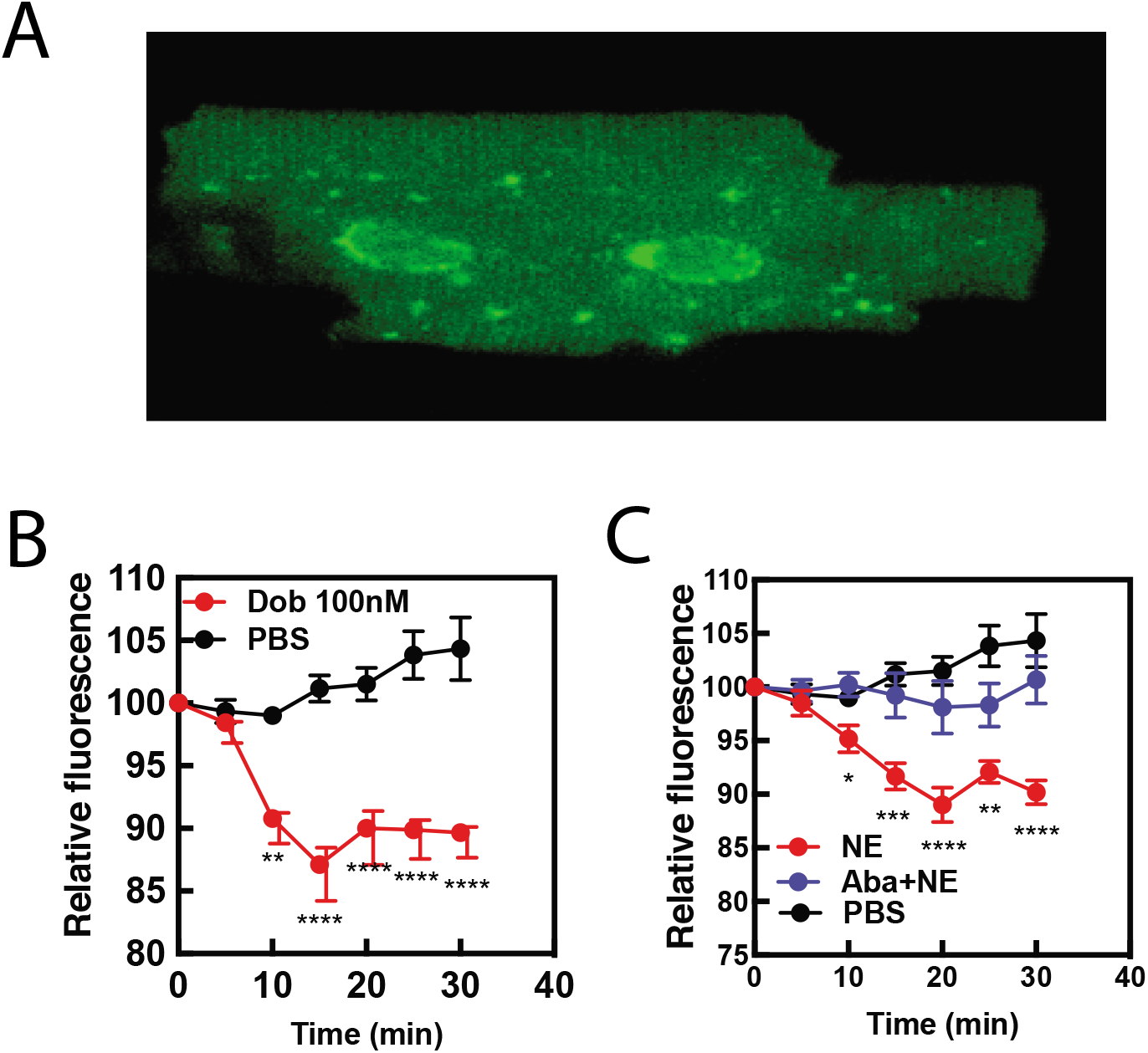
Dobutamine and NE stimulate PI4P hydrolysis in Adult Ventricular Myocytes (AVMs). Freshly isolated mouse AVMs were infected with FAPP-PH-GFP (50 MOI) for 24 hours. A) representative image of an AVM expressing FAPP-PH-GFP showing strong labeling surrounding the nucleus. B) FAPP-PH-GFP expressing AVMs were treated with either dob of PBS control as indicated and were imaged as for NRVMs as in Fig. 1A for PI4P hydrolysis. Fluorescence intensity in the region surrounding the nucleus was quantitated at 5 min intervals. C) AVMs were treated with PBS control, NE (10 μM) or NE plus abacavir and analyzed as in B. Data for B and C are from at least n=3 cells each from 3 separate preparations of AVMs.

### Cardiac hypertrophy induced by dobutamine is more effectively inhibited by a cell permeant antagonist

To determine if signaling by internal βARs is important for promotion of cellular hypertrophy, NRVMs were treated with dobutamine (100 nM) for 48 h and two measures of hypertrophy were assessed, cell area and ANF expression. Dobutamine stimulated an increase cell area (Fig. 7A), and ANF expression (Fig. 7B), after 48 hours of stimulation. Co-incubation with the cell permeant antagonist, metoprolol, strongly inhibited these hypertrophic responses. The cell impermeant antagonist, sotalol, on the other hand was significantly less effective at blocking these hypertrophy measures than metoprolol, (Fig. 7A and B), but did have a small effect on changes in cell area. These data, taken together with PI4P hydrolysis data, suggest that internal βARs are involved in mediating cardiomyocyte hypertrophy.

**Figure 7.**
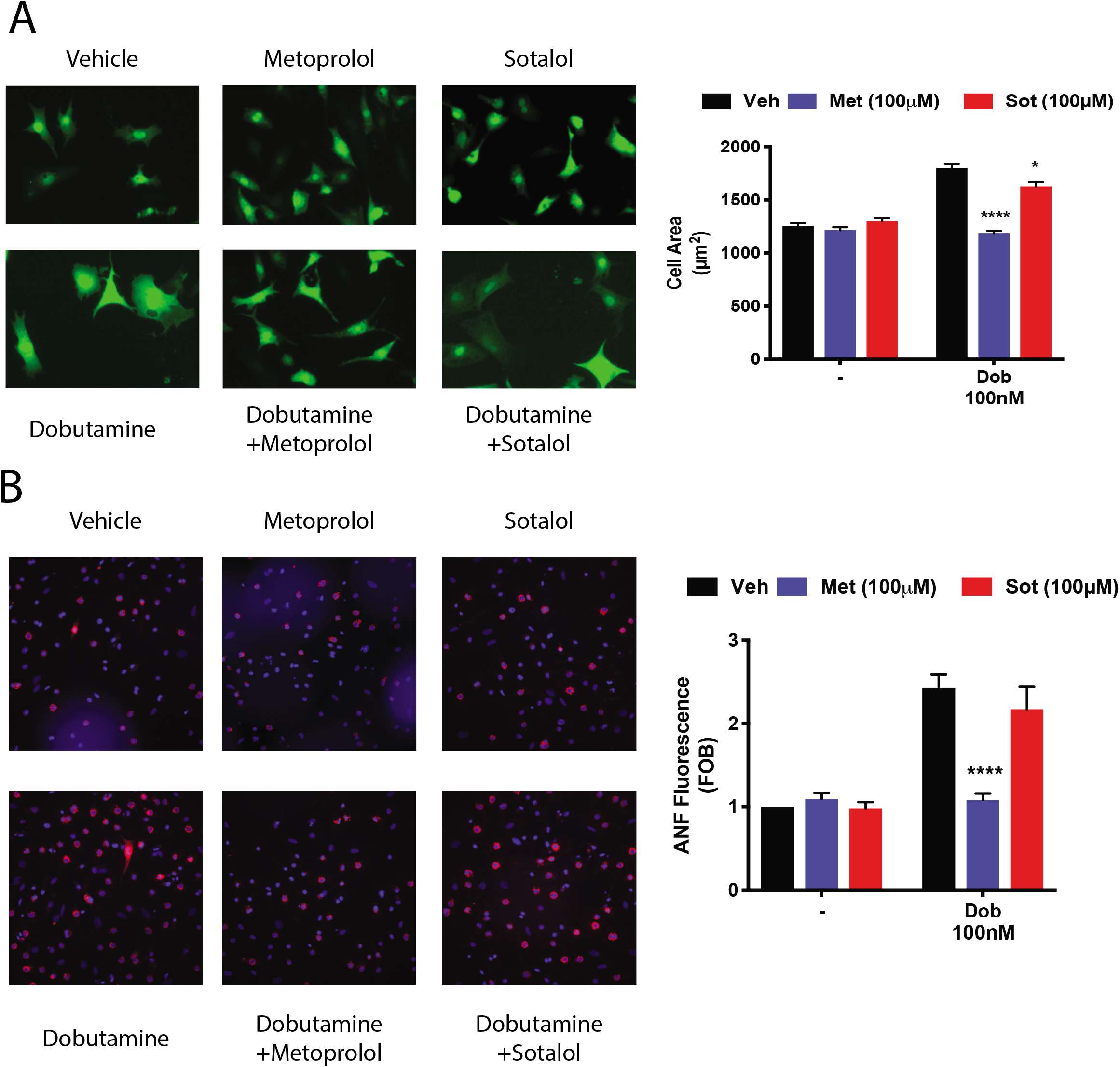
Dobutamine induced cardiomyocyte hypertrophy requires intracellular βARs. A) Dobutamine induces internal-receptor dependent increases in cell area. NRVMs were transduced with YFP virus prior to stimulation for 48 hours with dobutamine in the presence of the indicated antagonists or vehicle control. Following fixation, cell area was measured using image J. Representative images are on the left with mean data on the right. B) Dobutamine induces an increase in ANF expression via an internal receptor-dependent mechanism. NRVMs were stimulated with dobutamine in the presence or absence of the indicated antagonists or vehicle control for 48 hours before fixation and staining for ANF. Fluorescence of ANF rings was then captured by confocal microscopy, followed by fluorescence intensity analysis with Image J. Representative images are on the left with mean data on the right. All data is from at least 200 cells from 3 separate preparations of NRVMs.

### Golgi βARs and agonist internalization are required for stimulation of NRVM hypertrophy by NE

NE is an endogenously produced catecholamine involved in mediating hypertrophy and heart failure and stimulation of NRVMs with NE stimulates hypertrophy. To determine the role of intracellular NE in hypertrophy, NRVMs were treated with NE in the presence or absence of Corticosterone to block Oct3 cation transporters for 48h. Treatment with NE stimulated an increase in cell size (Fig. 8A) and ANF expression (Fig. 8B) that was blocked by Corticosterone indicating that transport of NE into the myocyte is required to mediate hypertrophy. Treatment with dobutamine also induced hypertrophy but this was not blocked by Oct3 supporting the idea that dobutamine access to internal receptors does not require Oct3 and confirms the specificity of the Oct3 blockade.

**Figure 8.**
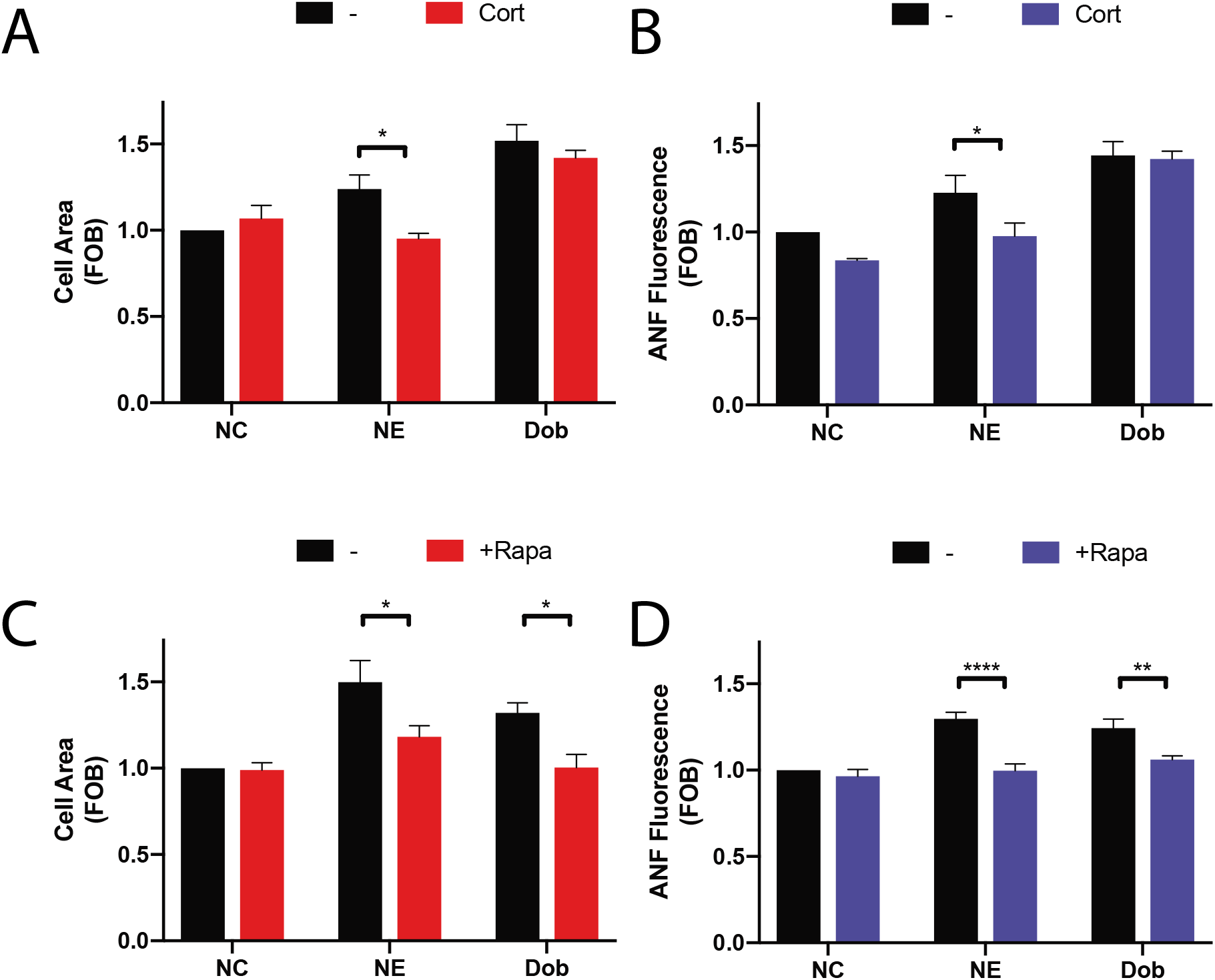
Dobutamine and norepinephrine induced cardiomyocyte hypertrophy requires Golgi-localized βARs. Norepinephrine induced hypertrophy also requires agonist internalization. Norepinephrine induces internal-receptor dependent increases in cell area (A) and ANF expression (B). NRVMs were stimulated for 48 hours with norepinephrine (10 μM) or dobutamine (100 nM) in the presence of Corticosterone (100μM) or vehicle control. Following fixation, cells were stained for ANF and using CellTracker Deep Red. Images were captured using a Thermo Fisher Cell Insight and analyzed by Cell Profiler. Both NE and Dobutamine significantly increase cell size (NE, p<0.05 and Dob p<0.001) and ANF expression (NE, p<0.01 and Dob p<0.001) in control conditions. Dobutamine and Norepinephrine require Golgi localized βAR to induce increase in cell area (A) and ANF expression (B). NRVMs were transduced with FRB-CFP-Nb80 and FKBP-mApple-GalT for 24 hours before stimulation. 15 minutes before agonist addition, Rapamycin or (1μM) or vehicle control was added. Cells were then stimulated with either dobutamine (100 nM) or norepinephrine (10 μM) for 48 hours. Following fixation, cells were stained for ANF and using CellTracker Deep Red. Images were captured using a Thermo Fisher Cell Insight and analyzed by Cell Profiler. Both NE and Dobutamine significantly increase cell size (NE, p<0.001 and Dob p<0.01) and ANF expression (NE, p<0.001 and Dob p<0.001) in control conditions. All data is from at least 1200 cells from 3 separate preparations of NRVMs.

To determine if NE and dobutamine driven NRVM hypertrophy requires Golgi βARs NRVMs were transduced with adenovirus containing CFP-Nb80-FRB and FKBP-mApple-GalT for 24 h followed by treatment for 48h with NE or dobutamine with and without co-treatment with rapamycin. In the absence of rapamycin, NE and dobutamine stimulated increases in cell area and ANF expression (Fig. 8C and D). Co-treatment with rapamycin to translocate Nb80 to the Golgi blocked NE and dobutamine stimulated hypertrophy, demonstrating that Golgi βARs are required for stimulation of hypertrophy by these agonists.

## Discussion

We previously found that raising cAMP levels with forskolin in NRVMs stimulated PI4P hydrolysis, but Iso did not unless a specific PDE, PDE3 was blocked. This led to the question of how the Epac/mAKAPβ/PLCε complex could be activated during physiological cAMP regulation by Gs coupled receptors. We considered two potential mechanisms, one where chronic agonist exposure alters the organization of PDEs in the cardiac myocyte such that Epac could be accessed by global cAMP changes, or that intracellular βARs generate cAMP with privileged access to the Epac/mAKAPβ/PLCε complex. Here we demonstrate that stimulation of Golgi resident β1ARs induces Epac-PLCε-dependent PI4P hydrolysis when stimulated by either a membrane permeant drug, dobutamine, or the physiological neurotransmitter, norepinephrine. We also show that stimulation of these internal receptors is required for both dobutamine and NE stimulated cardiomyocyte hypertrophy.

### Compartmentalized GPCR signaling

It is well established the GPCRs can be found on intracellular compartments (26). A number of investigators have characterized GPCRs on the nuclear envelope in neurons and cardiac myocytes, and signaling mechanisms have been proposed (27–30). GPCRs and G proteins have been found on other intracellular compartments including the Golgi apparatus, but signaling roles for receptors at the Golgi are not well defined. This is in part because GPCRs are trafficked through the Golgi apparatus on their way to the cell surface and it has not been clear that GPCRs in the Golgi have a signaling function. The recent work by Irranejad et al. (19) definitively demonstrated that Golgi localized receptors can indeed be driven to an active conformation by exogenous ligands. The work presented here elucidates a physiological function for these Golgi localized receptors through a definitive signal transduction pathway.

The purpose of internal GPCR signaling is likely to generate signals that are local and distinct from receptors at the PM or other intracellular locations. In this way specific processes can be controlled by second messengers that have the potential to activate a vast number of global responses if the signals were not restricted. In the case of the cardiac myocyte, during acute sympathetic stimulation, norepinephrine may only access cell surface βARs needed to mediate increases in cardiac contraction required to meet the demands of a sympathetic response. Because signaling by Golgi βARs requires uptake of ligand prior to activation these processes are slower and may not be accessed during acute sympathetic responses, but rather are only activated during more chronic exposure such as is observed in cardiovascular stress. Separation of the Epac/mAKAPβ/PLCε pathway at the nuclear envelope/Golgi interface from global cAMP dependent PKA activation likely prevents inappropriate activation of PLCε dependent hypertrophic responses to acute βAR activation.

### The requirement of internal receptors for cardiac hypertrophy

Significantly, stimulation of hypertrophy by the natural sympathetic ligand NE required activity of Golgi localized βARs. As has been discussed, sustained elevated sympathetic drive is one of the major factors responsible for development of cardiac hypertrophy due to chronic hypertension. β-blocker therapy efficacy is thought to result from preventing chronic stimulation of cell surface βARs by elevated catecholamines and subsequent desensitization (4). Our results suggest that blockade of internal receptors may be required to prevent catecholamine-induced hypertrophy and demonstrates that activation of cell surface GPCRs is not sufficient for natural catecholamines to induce hypertrophy in NRVMs. The importance of blockade of Golgi GPCRs is supported by the fact that the cell permeant βAR antagonist metoprolol is significantly more effective than the cell impermeant antagonist sotalol in prevention of dobutamine stimulated cardiomyocyte hypertrophy. Interestingly, clinically effective β-blockers tend to be relatively hydrophobic. Metoprolol and carvedilol are first line β blockers that are used to treat heart failure and have log P values 1.8 and 4.2 respectively, and are thus relatively cell permeant. Sotalol, which has a log P of −0.85 is relatively cell impermeant, and is not clinically used for heart failure treatment. We speculate that hydrophobicity might be an important factor for β-blocker development allowing these inhibitors to access internal pools of receptor.

A seeming contradiction is the observation that the cell impermeant agonist isoproterenol induces hypertrophy *in vitro* and when chronically administered *in vivo.* Iso administered *in vivo* is not at physiological concentrations and has multiple target tissues beyond the cardiac myocyte, such as the vasculature, that could contribute to hypertrophy. For NRVMs *in vitro*, while Iso can induce hypertrophy by a mechanism that probably does not rely on internal receptors, the effects of the natural ligands epinephrine and norepinephrine are more relevant to the *in vivo* situation and do rely on Golgi localized βARs. The synthetic agonist Iso seems to rely on an alternative PM mediated pathway that is not engaged by the physiological agonist NE. It will be critical to determine if in the *in vivo* pathology of heart failure that internal β receptors play a critical role using animal models such as TAC that mimic hypertensive stress.

## Materials and methods

### Isolation of neonatal cardiac myocytes and adenoviral transduction

Briefly, hearts were excised from 2 to 4 day old Sprague-Dawley rats, ventricles separated and minced thoroughly before digestion with Collagen type II (Worthington) in Hanks buffered saline solution (HBSS) without Ca^2+^ or Mg^2+^. Following digestion, cells were collected by centrifugation into Dulbecco Modified Eagle Medium (DMEM) supplemented with 10% fetal bovine serum (FBS), 100 U/mL penicillin, 100 μg/mL streptomycin, 2 mM glutamine and 2 μg/mL vitamin B12. Contaminating cells were removed by preplating cells onto tissue culture plastic for a minimum of 1 h at 37 °C. NRVMs were then plated onto either glass-bottom tissue culture plates or 12 well plates coated with 0.2% gelatin and cultured in DMEM (composition as above, with additional 10 μM cytosine arabinoside). 48 h later, cells were transferred into media supplemented with 1% FBS.

### NES-Venus-mini-Gs imaging

NRVMs were plated into gelatin-coated 20 mm glass bottom cell culture dishes. Cells were transfected the following day with plasmids (500-800 ng of β1-ARs and 250-400 ng of NES-Venus-mini-Gs per dish) using lipofectamine 3000. Media was changed to 1% FBS the next day and transduced with adenovirus-expressing CFP-Giantin overnight. Cells were imaged in confocal mode with a Leica DMi8 equipped with a Crest-optics X-light V2 confocal unit and a 100x 1.4 NA oil-immersion lens. Venus was excited at 515 with an X-Cite Xled1 light source, and emission monitored imaged on a backlit CMOS Photometrics Prime 95B camera.

### Measurement of PI4P hydrolysis

Measurements of PI4P hydrolysis were made as previously described (9, 11, 17). After preparation and culture of myocytes, cells were transduced with adenovirus (50 MOI) expressing GFP-FAPP-PH overnight. The following day, expression was confirmed by epifluorescence microscopy. Time lapse video fluorescence Imaging of GFP-FAPP-PH fluorescence was performed at room temperature or 37°C, where indicated, on a LEICA DMi8 with a 20x air lens in confocal mode. EGFP was excited at 488 nm with an X-Cite Xled1 light source and emission monitored imaged on a backlit CMOS Photometrics Prime 95B camera. Images were acquired with 50 ms exposure times at 1 min intervals to minimize photobleaching. Analysis of fluorescence intensity changes from the videos was performed using NIH Image J unless otherwise stated. Analysis was performed by subtracting background fluorescence intensity from a region of interest intensity at all time points measured. Data is presented as percentage of fluorescence remaining after agonist stimulation when compared to cells prior to stimulation.

### Measurement of myocyte cell size

NRVMs were plated into gelatin-coated 12 well plates or glass-bottom 96 well plates and allowed to grow overnight. The following day, cells were infected overnight with adenovirus expressing YFP. Subsequently, NRVMs were stimulated with dobutamine (100 nM) or norepinephrine (10μM) for 48 h, in the presence of antagonists or transduction with FRB-CFP-Nb80 and FKBP-mApple-GalT, as indicated. Following stimulation, cells were fixed in 4% (w/v) paraformaldehyde. Fluorescent images were taken at 10 x magnification and cell area measured using NIH Image J or Cell Profiler software from over 500 cells from at least 3 separate experiments.

### Immunocytochemistry for ANF induction

NRVMs were plated into gelatin-coated 8 chamber glass slides or glass-bottom 96 well plates and allowed to grow overnight. The following day, cells were serum starved for 24 hours. Subsequently, NRVMs were stimulated with dobutamine (100 nM) or norepinephrine (10μM) for 48 h, in the presence of antagonists or transduction with FRB-CFP-Nb80 and FKBP-mApple-GalT, as indicated. Cells were washed with PBS and fixed with 4% PFA for 15 minutes and then incubated with 10% normal goat serum in phosphate buffered saline containing 0.1% Triton X100 (PBS-T) for 1 hour at room temperature. Primary antibody was incubated at a dilution of 1:1000 in 2 % goat serum in PBS-T overnight at 4C. After three washes with PBST, cells were incubated with secondary antibody (Goat anti-rabbit Alexa Fluor 568) at a dilution of 1:1000 in PBS-T for 1.5 hours at room temperature. After three washes with PBS-T, DAPI was added at a dilution of 1:500 in PBS and incubated for 30 minutes. Fluorescence images were captured at 10 x magnification and fluorescence intensity corresponding to ANF staining surrounding the nucleus was quantified using either NIH Image J or Cell Profiler software.

### Inhibition of Golgi βARs by Nb80

NRVMs were plated onto 20 mm glass bottom coverslips and allowed to adhere for 24 hours. Following adhesion, cells were transduced with adenoviruses containing FKBP-mApple-GalT construct (a kind gift from R. Irannejad, UCSF) and FRB-CFP-Nb80 (modified from R. Irannejad, UCSF). The following day, NRVMs were transduced with adenovirus (50 MOI) expressing GFP-FAPP-PH overnight. 15 minutes prior to experimentation, cells were treated with either 1 μM rapamycin or DMSO, where indicated, to induce translocation of Nb80 to the Golgi membrane. Imaging of GFP-FAPP-PH fluorescence was performed at 37°C on a LEICA DMi8 in confocal mode with a 20 x air lens. Analysis was performed as for PI4P hydrolysis as previously indicated.

#### Statistical Analysis

All graphs are presented as mean ± SE. Agonist treatments were compared to vehicle control performed on the same day and were added where indicated by the arrow. All data was analyzed by two-way unpaired ANOVA with Sidak’s post-hoc test. * p<0.05 ** p<0.001 *** p<0.0001 **** p<0.00001 using GraphPad Prism 7.0.

## Acknowledgements

Supported by NIH Grant R35GM127303

## This PDF file includes

Fig S1

Legends for movies S1 to S5

## Other supplementary materials for this manuscript include the following

Movies S1 to S5

**Fig. S1.**
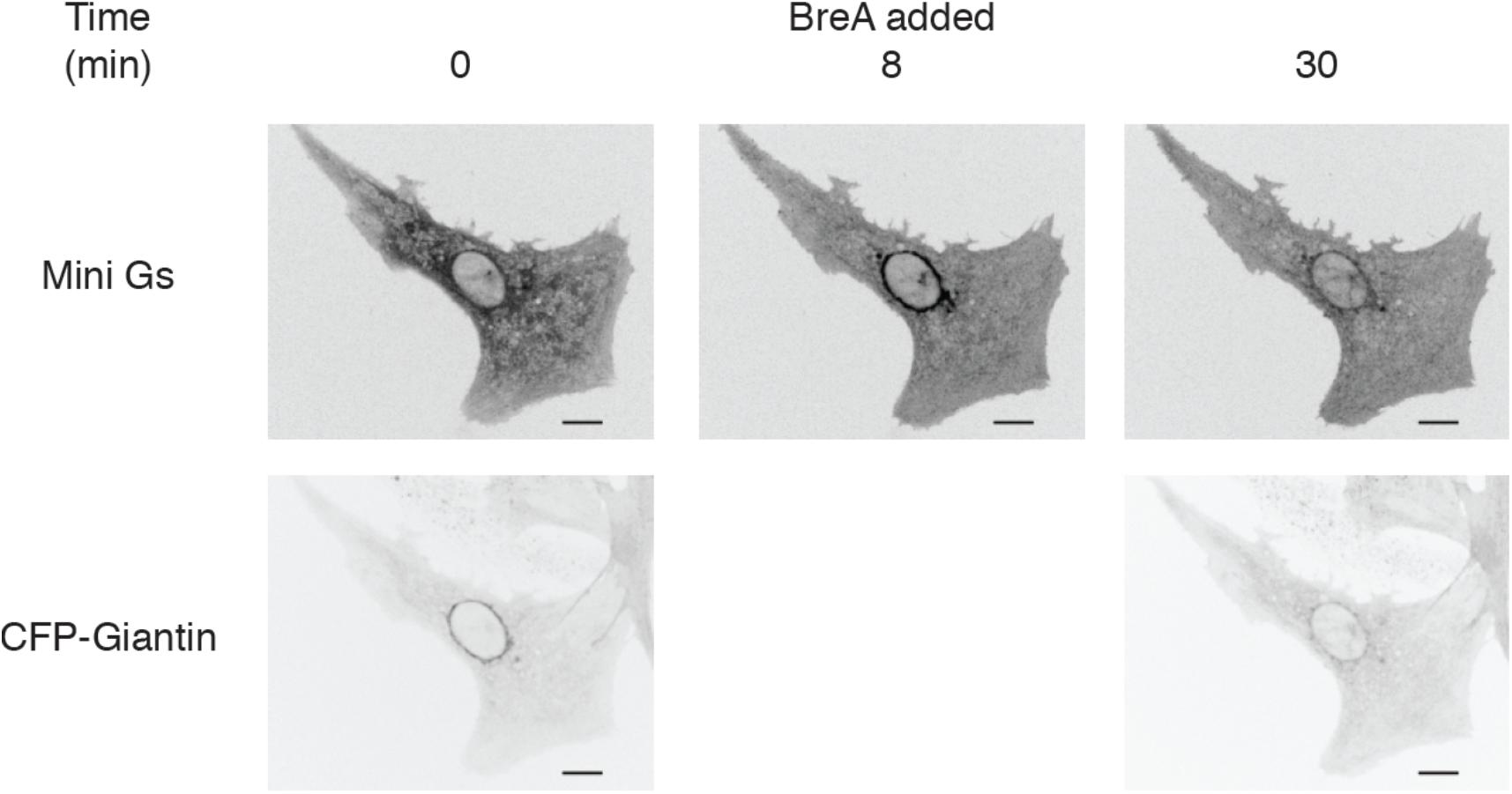
Disruption of the Golgi apparatus reverses mini-Gs protein recruitment to the perinuclear region by dobutamine. NRVMs were transfected with β1-AR and NES-Venus-mini-Gs, followed by viral transduction with CFP-Giantin. Representative images of dobutamine-mediated NES-Venus-mini-Gs recruitment (100 nM, Upper panels) and CFP-Giantin Golgi marker (lower panels). Dobutamine was added at 2 min and Brefeldin A (5μg/ml), was added at 8 min just after the 8 min image above was captured. NES-Venus-mini-Gs recruitment continually monitored.

Movie S1. Dobutamine activates β1ARs in the Golgi apparatus. Confocal images of β1-ARs-overexpressing cardiomyocytes with NES-Venus-mini-Gs, treated with 100 nM dobutamine. Total time represented by the movie is 10 min and pictures were taken every 15 seconds. Dobutamine was added at 2 min timepoint.

Movie S2. Isoproterenol does not activate β1ARs in the Golgi apparatus. Confocal images of β1-ARs-overexpressing cardiomyocytes with NES-Venus-mini-Gs, treated with 100 nM isoproterenol. Total time represented by the movie is 10 min and pictures were taken every 15 seconds. Isoproterenol was added at 2 min timepoint.

Movie S3. Disruption of the Golgi apparatus inhibits activation of β1ARs in the Golgi apparatus. Reversal of mini Gs recruitment to Golgi membrane after addition of Brefeldin A. Total time represented by the movie is 30 min and pictures were taken every 30 seconds. Dobutamine was added at 2 min timepoint for 8 min and then Brefeldin A was added for next 20 min.

Movie S4. Norepinephrine activates β1ARs in the Golgi apparatus. Confocal images of β1-ARs-overexpressing cardiomyocytes with NES-Venus-mini-Gs, treated with 10 μM norepinephrine. Total time represented by the movie is 30 min and pictures were taken every 30 seconds. Norepinephrine was added at 2 min timepoint.

Movie S5. Inhibition of receptor internalization does not alter activation of β1ARs in the Golgi apparatus. Confocal images of β1-ARs-overexpressing cardiomyocytes with NES-Venus-mini-Gs, pretreated with 40 μM Dyngo. Total time represented by the movie is 30 min and pictures were taken every 30 seconds. Dyngo was added 15 min prior to experimentation and norepinephrine was added at 2 min timepoint.

